# Neurocognitive consequences of hand augmentation

**DOI:** 10.1101/2020.06.16.151944

**Authors:** Paulina Kieliba, Danielle Clode, Roni O Maimon-Mor, Tamar R. Makin

## Abstract

From hand tools to cyborgs, humans have long been fascinated by the opportunities afforded by augmenting ourselves. Here, we studied how motor augmentation with an extra robotic thumb (the Third Thumb) impacts the biological hand representation in the brains of able-bodied people. Participants were tested on a variety of behavioural and neuroimaging tests designed to interrogate the augmented hand’s representation before and after 5-days of semi-intensive training. Training improved the Thumb’s motor control, dexterity and hand-robot coordination, even when cognitive load was increased or when vision was occluded, and resulted in increased sense of embodiment over the robotic Thumb. Thumb usage also weakened natural kinematic hand synergies. Importantly, brain decoding of the augmented hand’s motor representation demonstrated mild collapsing of the canonical hand structure following training, suggesting that motor augmentation may disrupt the biological hand representation. Together, our findings unveil critical neurocognitive considerations for designing human body augmentation.

## Introduction

There are many things that we could do better if we had more fingers in our hand. Engineers are currently developing extra robotic fingers and even entire arms aimed to augment our bodies by expanding our natural motor repertoire (*1–5*). Despite rapid advancements in augmentative technologies, little notice is given to the crucial question of how the human brain might support them. The augmentative devices aim to change the way we interact with the environment, which entails changes to how we move and operate our biological body. Here we asked whether the human brain can accommodate motor control of an extra robotic finger, focusing on its impact on the neural representation of the biological hand.

Hand representation in the primary sensorimotor cortex of the brain has a well-established functional structure that develops very early on (*6, 7*). It is highly consistent within (*8*) and across (*9*) participants and is preserved even after severe loss of motor functions due to e.g. stroke (*9*), spinal cord injury (*10*), disability (*11*) or even hand amputation (*12–14*). Hand representation has been suggested to reflect daily hand use (*9*), with studies showing that it may be altered under constrained circumstances. Most notably in musicians’ dystonia, a clinical condition involving increased finger enslavement, the individualised representation of single fingers has been shown to collapse (*15*, though see *16*).

Here we trained able-bodied people to use an extra robotic thumb (the Third Thumb, created by Dani Clode (*17*), hereafter “Thumb”) over the course of 5 days, including both lab-based and in-the-wild daily use. The Thumb is a 3D-printed supernumerary robotic finger, with two degrees of freedom, controlled with pressure exerted with the big toes, designed to extend the natural repertoire of hand movements (Figure 1A-B). During training, we tracked (biological) finger coordination and compared it with normal hand use. We tested for changes in motor control and embodiment of the Thumb, as well as hand-Thumb coordination before and after training. Augmented participants were compared to a control group that underwent similar training regime while wearing the Thumb without being able to control it. We also examined how the sensorimotor and body representation of the augmented hand changed following Thumb training. We hypothesised that successful hand-robot cooperation and subsequent change of finger co-use will update both biological and artificial body representation.

**Figure 1.**
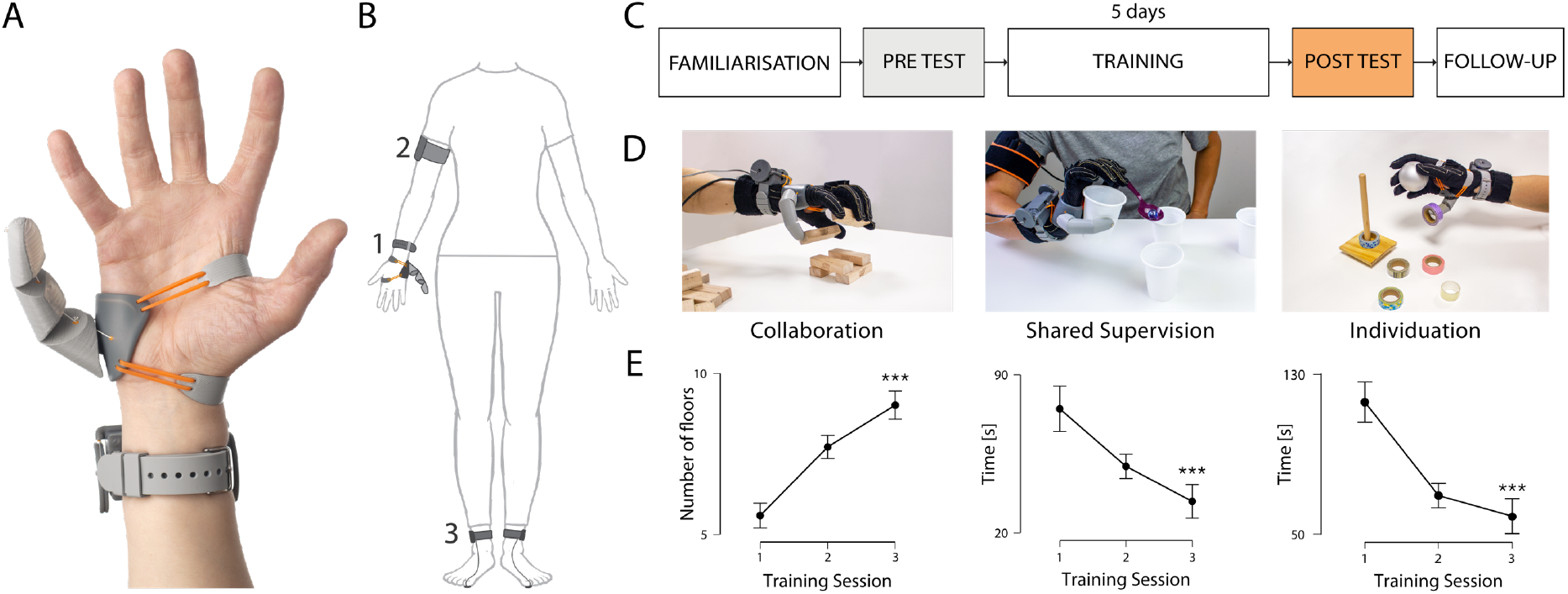
Experimental design. (A-B) The Third Thumb is a 3D-printed robotic thumb. Mounted on the side of the palm (1), the Thumb is actuated by two motors (fixed to a wrist band), allowing for independent control over flexion/abduction. The Thumb is powered by (2) an external battery, strapped around the arm and wirelessly controlled by (3) two force sensors strapped underneath the participant’s big toes. (C) Experimental design for the augmentation group. (D-E) Examples of the inlab training tasks used for hand-Thumb collaboration (building a Jenga tower), shared supervision (holding a plastic cup while extracting a marble with a spoon) and Thumb individuation (stacking tapes with the Thumb while the biological hand is occupied). Participants showed significant performance improvements across training session. Asterisks denote significant effect of time at *** p<0.001.

## Results

### Improved motor control, hand-Thumb coordination and Thumb embodiment following daily Thumb usage

We first characterised motor performance of the augmented hand throughout the 5 days of usage. Augmentation participants completed five daily in-lab training sessions (1.58±0.22hr; mean±std) and were additionally encouraged to use the Thumb outside the lab (2.61±1.18hr; self-reported). The average use time, as quantified by the automatic usage logs, was 2.95±0.84hr per day, out of which a total of 1.37±0.49hr involved active Thumb movement.

During daily training sessions, participants were presented with a variety of reaching, grasping and in-hand manipulation tasks designed to create different challenges for the Thumb usage (see Supplementary Video 1 for examples). Augmentation participants showed significant improvement on all the training tasks (main effect of time for all tasks: p<0.001, η_p_^2^>0.5 Figure 1D-E & Supplementary Figure S1). Motor control was further assessed using a hand-Thumb coordination task, requiring participants to oppose the Thumb to their biological fingers. Even though controlling the Thumb with the big toes may seem unusual, participants were able to successfully perform the hand-Thumb coordination task even at baseline (Figure 2B), though this performance was significantly improved after training. Significant improvements were observed both during daily training (F(4,76)=28.24, p<0.001, η_p_^2^=0.6 Figure 2A-C), and when comparing the performance pre- and post-the 5 days of training using a sequential variation of the same task (see Methods). Here, augmentation participants showed significant improvements, not only with vision (t(19)=8.96, p<0.001, η_p_^2^=0.81), but also when blindfolded (t(19)=7.40, p<0.001, η_p_^2^=0.74, Figure 3A), indicating improved Thumb proprioception. Importantly, the participant’s motor performance with the Thumb was not impacted by increased cognitive load incurred during a dual-task involving numerical cognition and motor control, performed during the first and last days of the training. This was demonstrated by the lack of significant cognitive load x session interaction (F(1,16)=0.003, p=0.959) and a non-significant main effect of the cognitive load (F(1,16)=2.465, p=0.136; Figure 2D). This was further confirmed using a one-tailed Bayesian t-tests, which yielded a Bayesian Factor (BF) of 0.13 and 0.14 for the first/last day of training, indicating moderate evidence in support of the null hypothesis. As such, participants learned to operate the Thumb under a variety of circumstances, extending beyond their specific training.

**Figure 2.**
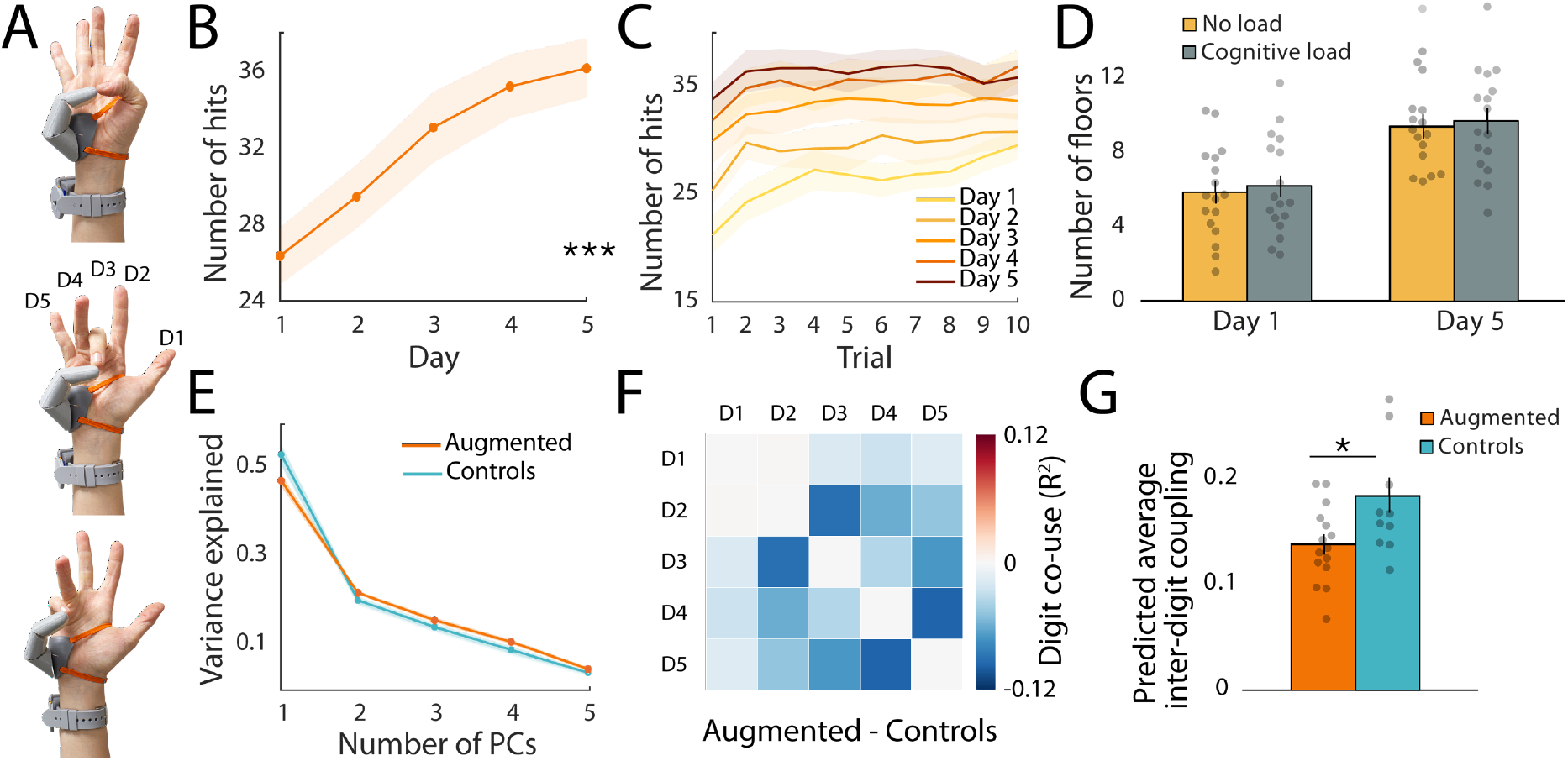
Training outcomes. (A-C) Augmentation participants showed significant daily improvement on the hand-Thumb coordination task. (D) Motor performance with the Thumb was not impacted by increased cognitive load during the first and last training days. (E) Hand kinematics data collected during the training sessions. The first principal competent (synchronised movement across all five fingers) captured less variance in the augmentation group compared to controls, indicating less synchronised movements. (F-G) The augmentation group showed lower inter-finger coupling, relative to controls during Thumb use, indicating change to the natural finger coordination. The bars depict group means, error bars represent standard error of the mean. Individual dots correspond to individual subjects’ average inter-finger (D1-D5) coordination scores as predicted by the linear mixed model (see Methods). Asterisks denote significant effects at * p<0.05 and *** p<0.001.

**Figure 3.**
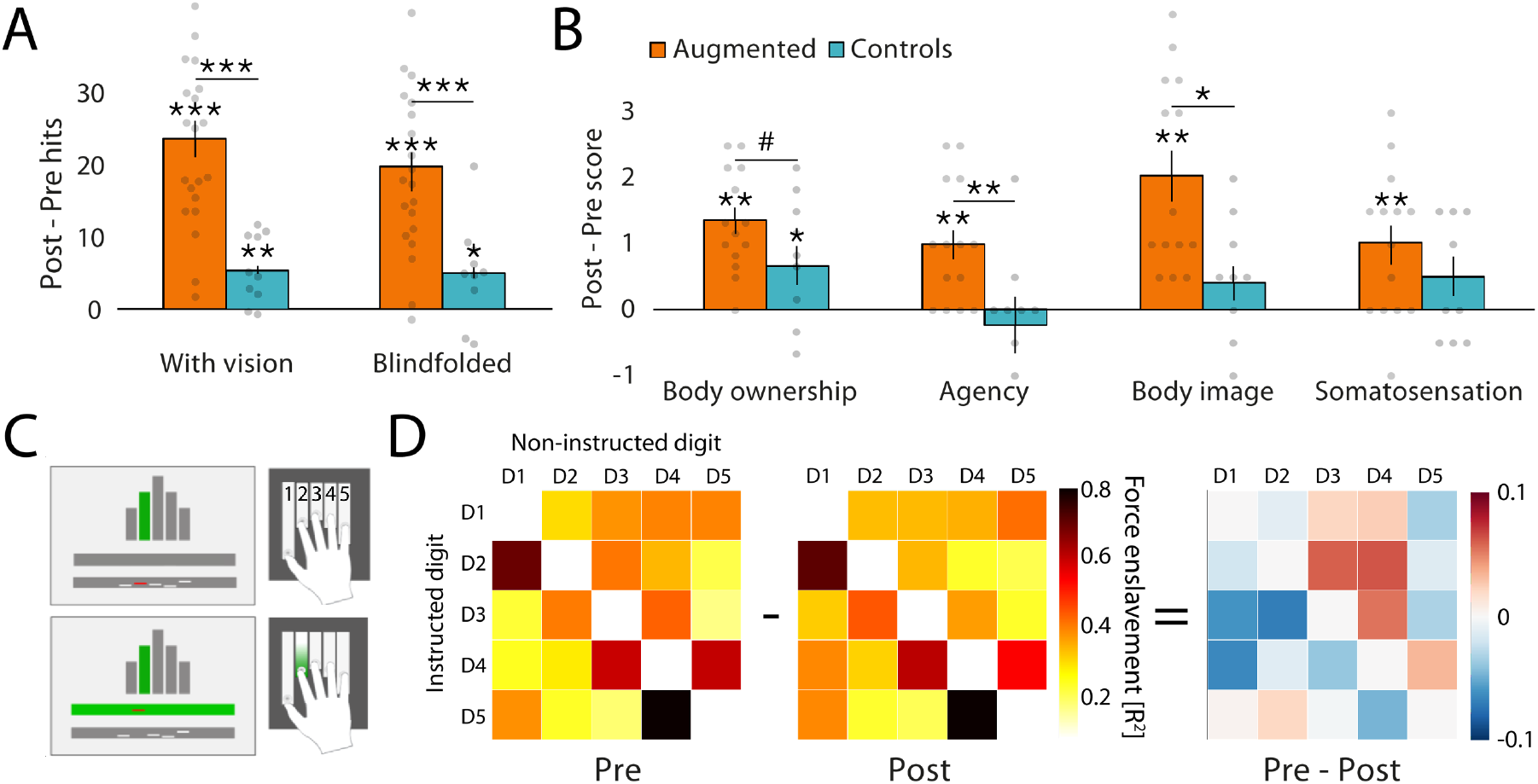
Behavioural correlates of hand augmentation. (A) Augmentation participants showed greater improvement than controls on a hand-Thumb coordination task conducted before and after the training period. Participants showed improved performance even while blindfolded, indicating increased Thumb proprioception. (B) Self-reported Thumb embodiment increased significantly in the augmentation group following Thumb training. (C-D) Participants performed individuated key presses with an instructed finger, while the forces exerted with the non-instructed fingers were measured to obtain a 5×5 force enslavement matrices. Matrices presented here depict the group average. No significant change in the average enslavement was observed. Asterisks denote significant effects at * p<0.05, ** p<0.01, *** p<0.001.

We also assessed the perceived sense of embodiment over the Thumb following the training period, relative to baseline. Participants were asked to respond to statements relating to key embodiment features (*18, 19*). Augmentation participants reported a significant increase of embodiment in each of the four categories (body ownership: t(13)=6.57, p<0.001, η_p_^2^=0.77; agency: t(13)=4.07, p<0.001, η_p_^2^=0.56; body image: t(13)=5.215, p<0.001, η_p_^2^=0.68; somatosensation: t(13)=6.032 p<0.001, η_p_^2^=0.74; Figure 3B).

As the improvements described above could be skewed due to task repetition, we also tested a group of 11 control participants, who underwent similar pre- and post- tests and training regime (see Methods) but wore a static version of the Thumb for the same duration of time - 4.11±1.06hr per day (t(29)=0.526, p=0.6, BF=0.39), out of which 2.93±1.34 outside of the lab-based training sessions (t(29)=-0.697, p=0.49, BF=0.42 for wear-time group comparison). The control group showed significantly lower improvement on the pre-post sequential hand-Thumb coordination (significant effect of group revealed by ANCOVA - with vision: F(1,27)=22.86, p<0.001, η_p_^2^=0.44; blindfolded: F(1,27)=11.96, p=0.002, η_p_^2^=0.28) and did not embody the extra Thumb to the same extent as the augmentation group (agency: F(1,21)=10.013, p=0.009, η_p_^2^=0.285; body image: F(1,21)=11.16, p=0.012, η_p_^2^=0.26). For body ownership, the group effect was trending (F(1,21)=4.07, p=0.057, η_p_^2^=0.16), while for somatosensation scores, the group comparison was nonsignificant (BF=0.48). These results indicate that active usage is critical for developing proprioception and embodiment of the robotic Thumb.

### Hand augmentation impacts motor control of the natural hand

Next, we investigated the impact of hand augmentation on motor control of the augmented (right) hand. We first examined the complexity of the hand movements (i.e. kinematic synergies) during Thumb use, captured with a Cyberglove during the in-lab training. We found that in general more principal components (kinematic synergies) were needed in the augmentation group, compared to the control group, to explain the 80% of the total variance of the hand movements (F(1,22)=5.52, p=0.03, η_p_^2^=0.2, Supplementary Figure S3B). This difference was however strongly driven by the amount of variance explained by the first principal component, corresponding to the coordinated flexion of all fingers (Supplementary Figure S3A). Indeed, the variance explained by this inter-finger synergy was significantly decreased in the augmentation group compared to controls (F(1,22)=6.27, p=0.02, η_p_^2^=0.22, Figure 2E), while no difference was found between the first and the last day of training (F(1,22)=2.57, p=0.12, BF=0.62). Since the remaining principal components represent more intricate finger movements, the decrease of variance explained by the first kinematic synergy suggests more finger individuation in the augmentation group. To uncover more detailed changes in finger coordination, we then looked at the degree of kinematic coupling between individual digit pairs. Here again, no differences in finger coordination were found between the first and the last day of training (main effect of time: F(1,23)=1.3, p=0.27, BF=0.17). For the augmentation group, this finding indicates that the strategies implemented for incorporating the Thumb into the motor repertoire during day 1 were generally preserved throughout training. Consistent with the PCA results, we found significant differences in finger coordination implemented across groups (group x finger-pair interaction: F(9,414)=2.66, p=0.005), with an overall decrease in inter-finger coupling in the augmentation group relative to controls (main effect of group: F(1,23)=6.98, p=0.01, Figures 2F-G). Together these results indicate a potential breakage of natural finger synergies during Thumb use.

Changes in inter-finger motor control were further investigated through force enslavement (involuntary force production by non-intended fingers), measured pre- and post- Thumb use (Figure 3C-D). While no significant changes to the overall pattern of the force enslavement were found, in the augmentation group there was a trend towards increased enslavement caused by the biological thumb following extra Thumb use (F(1,17)=3.36, p=0.08, first column of the pre-post difference matrix in Fig3D). However, this effect was not robust, as demonstrated by a lack of significant time x group interaction when comparing the augmentation group with controls (F(1,27)=1.5, p=0.23, BF=0.57).

We also examined potential changes to body image (perceptions and attitudes concerning one’s body representation; (*20*)). Using a tactile distances task, we found no significant pre- to post- changes in tactile biases (t(18)=0.164, p=0.87, BF=0.24). Similarly, we did not observe any significant changes incurred by Thumb usage using a visual hand lateralisation task. While participants did get faster at visual hand lateralisation (F(1,16)=6.89, p=0.018), this was the case for judgements made both for the augmented and the non-augmented hand (hand x session interaction F(1,16)=0.019, p=0.89, BF=0.326). These findings indicate that hand augmentation does not influence all aspects of body representation, with body image remaining stable irrespective of Thumb usage.

### Canonical representation structure shrinks following extra Thumb use

Having observed that augmentation participants show changed pattern of finger coordination when compared to the controls, we sought to understand whether Thumb usage can alter the biological hand representation in the sensorimotor cortex. As such, we used fMRI to compare neural hand representation before and after Thumb use. During the scans, participants were required to make individuated finger movements with their biological fingers. Note that due to MRI safety considerations, participants were not wearing the Thumb during the scans. Here we also tested the non-augmented (left) hand, providing a within-participant control.

First, we used a univariate approach to interrogate activity levels within the (independently localised) cortical territory of the biological hands. We found that the average activity was decreasing from the pre- to post- scan (main effect of time: F(1,18)=7.89, p=0.012, η_p_^2^=0.3). However, this decrease was not specific to any of the hands (hand x time interaction: F(1,18)=1.66, p=0.21, BF=0.37).

The bars depict group mean, error bars represent standard error of the mean. Individual dots correspond to individual subjects’ average distance as predicted by the linear mixed model (see Methods). (D) Multidimensional scaling (MDS) depiction of the left and right (augmented) hand representational structures. Ellipses indicate between-participant standard errors. Darker colours represent the post scan, whereas lighter colours represent the pre (baseline) scan. Red = D1, Yellow = D2, Green = D3, Blue = D4, Purple = D5. Asterisks denote significant effects at * p<0.05 and *** p<0.001.

To investigate changes to the augmented hand’s representational structure, we estimated the dissimilarity between activity patterns elicited by individual fingers’ movements in the sensorimotor cortex, as measured using cross-validated Mahalanobis distance (*21*). Small inter-finger dissimilarity values indicate that the representation of the two fingers is more similar/overlapping, while larger dissimilarity implies more individuated finger representation. Augmentation participants showed significantly reduced dissimilarity of the augmented (right) hand representation in the sensorimotor cortex following Thumb use (t(25.1)=2.3, p=0.03, Figure 4). This shrinking effect was specific to the augmented hand, as demonstrated by a significant hand x time interaction (F(1,722)=12.89, p<0.001, Figure 4).

**Figure 4.**
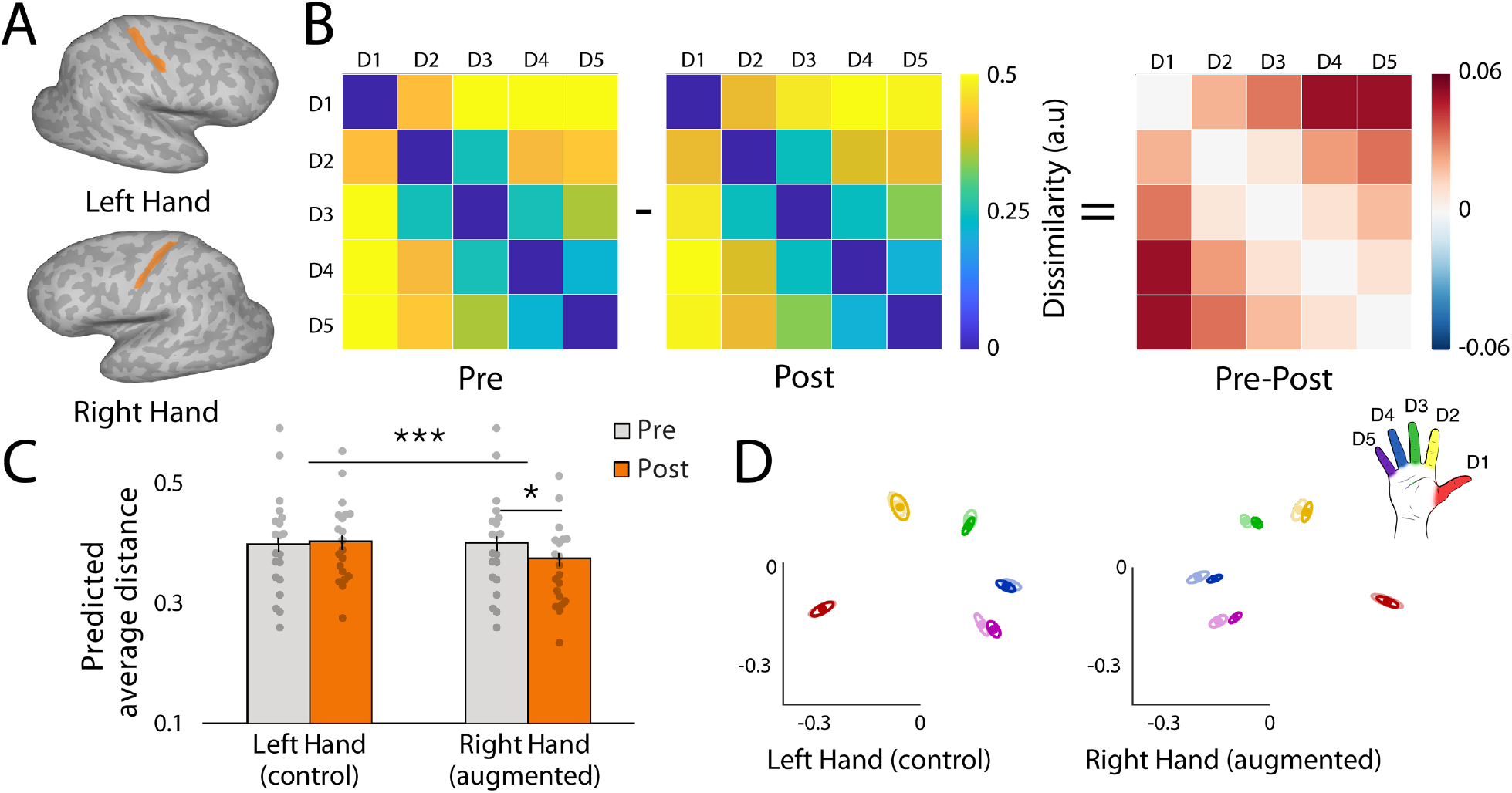
Canonical hand representation shrinks following hand augmentation. (A) The sensorimotor ROI was defined using anatomically defined M1 hand ROI including Broadmann area BA4 (Freesurfer segmentation), partially overlapping the central sulcus. (B) Group mean dissimilarity matrix of the right (augmented) hand pre- and post- training. (C) The average inter-finger dissimilarity of the right (augmented), but not the left (non-augmented) hand decreased significantly following Thumb use.

The reduction of inter-finger dissimilarity was diminished in a follow-up scan, conducted 7-10 days after the end of training, collected from a subset of available participants (n=12). This was supported by moderate evidence (BF=0.31) for a null difference between the average distances measured during pre- and follow-up sessions, indicating that the shrinkage of the canonical hand representational structure depends on recent Thumb use (note however that with this subset of participants the initial difference between the pre and post sessions was ambiguous – BF=0.58). We also examined the inter-finger representational structure’s typicality, that is the correlation of the individual’s representational dissimilarity matrix (RDM) with a group average RDM, computed from the pre-data, and found no significant pre- to post- differences (BF=0.67). These findings show that using the extra Thumb not only alters the motor control of the biological hand, but also impact the representational similarity of individual fingers’ representation in the brain. Crucially, this effect was observed while participants were not using or even wearing the Thumb.

### Effective connectivity between hand and feet motor areas remains invariant

Finally, we explored the relationship between the hand’s and toes’ (controlling the Thumb’s movements) representation in the sensorimotor cortex, but found no significant differences following training, relative to baseline. First, we used a univariate approach to examine feet-specific activity within the augmented hand territory, and found that it had not increased significantly (t(18)=0.809, p=0.43, BF=0.27). Next, we examined the functional coupling between the sensorimotor hand and feet areas, using resting-state functional connectivity analysis. We extracted the mean time course from the augmented and non-augmented hands’ territories, as well as from the feet territory, and used partial correlations to investigate the resting-state coupling between the augmented hand and the feet (while accounting for connectivity between the two hands; t(19)=0.774, p=0.45, BF=0.3). Together, these results suggest that while augmentation might promote plasticity locally (i.e. between fingers), the global homunculus likely remains largely unchanged.

## Discussion

Here, we provide the first comprehensive demonstration of successful motor integration of a robotic augmentation device (the Third Thumb) and explore how augmentation impacts the user’s sensorimotor hand representation in the brain. After only 5 days of Thumb use, participants showed significant improvements in six-fingered motor performance across multiple tasks. In addition to individuated control of the extra Thumb, participants were able to integrate Thumb motor control with the movements of their natural hand, requiring collaboration, shared supervision and hand-robot coordination. Motor performance was greatly improved even without visual feedback and remained stable under increased cognitive load. We further show that hand augmentation resulted in increased sense of embodiment over the Thumb - a key goal for successful augmentation (*22*). By demonstrating successful adaptation to motor augmentation under diverse ecological task demands, our findings constitute a leap beyond earlier pioneering proof-of-concept accounts of successful usage of extra robotic fingers (*1, 4, 23–25*) or arms (*3, 5*) under restricted circumstances.

Importantly, successful adoption of augmentative technologies relies not only on the user’s proficiency in operating the robotic device. A further challenge for augmentation is to ensure that the device usage will not impact users’ ability to control their biological body, especially while the augmentative device is not being used. Therefore, a critical question for anyone interested in safe motor augmentation is whether it would incur any costs to the user’s biological body representation. This concern is rooted in previous research of brain plasticity, demonstrating that our motor experience shapes the structure and function of the nervous system (*9*). As such, since motor augmentation is designed to change the way we interact with the environment, it is reasonable to predict that it will reshape the neural basis of our biological body. With that in mind, our investigation was focused on changes incurred to the biological body representation even while the Thumb isn’t being operated. This approach allows for our findings to be easily generalised to any other form of human-robot interfaces.

Traditionally, body representation in the sensorimotor cortex is considered to be highly adaptive even in the adult brain (*26, 27*) however recent research contributes a new perspective on its malleability (*12, 13*). Tools have been suggested to update the biological body representation, for example by tool-body integration (*28–30*). Yet, tools are normally used to replace the capacity of the hand, rather than to accompany it. Therefore, when using a tool, one is not required to radically alter their hand function (for example, the user will choose a grip for the tool’s handle that fits the natural synergies of the fingers). As such, tool-use does not entail an updated representation of the hand itself. Conversely, motor augmentation invites the user to reinvent the way they use their own body. This challenge is more closely akin to the acquisition of a new and complex motor skill – e.g. learning to play the pian0. Here, research has demonstrated that long-term training leads to change in finger representations (*31*). Specifically, trained pianists (over the course of many years) demonstrate reduced representational selectivity (lower representational dissimilarity) relative to novices. This evidence further emphasises the need to examine how long-term motor augmentation can impact the biological hand representation.

Here, we used a variety of pre- to post- measures to study changes in body representation following 5 days of Thumb usage. While some aspects of body representation (body image, large scale connectivity profile) were found to be stable, semi-intensive Thumb usage (2.3–6.3 hours per day) also resulted in mild, yet significant changes to the sensorimotor hand representation. First, motor integration of the Thumb resulted in breakage of the natural finger coordination patterns (kinematic synergies) during Thumb use, previously considered as a key determinate of sensorimotor brain organisation (*9, 32, 33*). Such an abrupt change in finger coordination may ‘disorganise’ the existing hand representation. Evidence for altered sensorimotor control of the biological hand was also apparent after training, when the Thumb was not being used, or even worn. Specifically, we observed a reduction of representational selectivity (i.e. dissimilarity) between representational patterns of the 5 fingers, akin to increased inter-finger overlap which had been previously used as a marker for decreased motor control (*15*). This effect may be reversible (as indicated by the follow-up scan taken 7-10 days after Thumb usage had ceased), or even transient. Yet, this evidence nevertheless suggest that motor augmentation might incur some costs to the augmented hand representation. It is therefore crucial to consider what plasticity mechanisms might be triggered during Thumb use to cause the observed changes in the biological hand representation.

The “maladaptive” plasticity account highlights the fact that neural resources devoted to hand representation are limited, and thus changes to the existing hand representation might incur a cost on motor control (*15*). Here, the adoption of an extra robotic thumb by an adult with a stable hand representation will promote a change to existing brain organisation. We used computational simulations to explore the feasibility of multiple potential plasticity mechanisms that could trigger the observed fMRI results (reduction in pairwise distances between fingers; Supplementary Figure S2). First, based on information processing theories (*34, 35*), the integration of a new additional finger into the hand’s motor control could impinge on the existing representation of the biological fingers. Second, the change in finger coordination, observed during training, may also lead to abrupt changes in excitability profiles that can trigger homeostatic plasticity mechanisms and promote increased tonic, and relatively wide-spread inhibition (*36*). Thirdly, changes to finger coordination during Thumb use can result in increased finger individuation, leading to increased cortical representation of individual fingers via Hebbian learning (cortical magnification, (*37*)). By simulating “neuronal activity” over a fixed-size ROI split into finger specific areas (see Methods), we found that each of these processes is conceptually capable of causing the observed reduction in representational selectivity (Supplementary Figure S2). It is also possible that these distinct mechanisms interact with each other as they may all be relevant to the experience of using robotic motor augmentation. This is in stark contrast to a recent report of individuals who were born with a 6^th^ (fully operational) finger and could harness processes of developmental plasticity to establish normal motor control across all six fingers (*38*). Finally, it is worth considering adaptive, rather than maladaptive plasticity processes that could be involved in developing motor control over a new body part. For instance, by inducing new kinematic synergies, learning to use the extra Thumb may be pushing the network outside of its existing manifold (*39*), to allow for formation of new neural activity patterns directed at optimal representation of the Thumb relative to the rest of the augmented hand.

To conclude, emerging technologies, designed to assist, substitute and even augment our motor abilities hold tremendous promise for transforming the lives of both disabled and healthy communities. This vision depends not only on the exciting technological innovations, it also critically relies on our brain’s ability to learn, adapt and interface with these devices. Therefore, as technology becomes more integrated with the human body, we see new challenges and opportunities emerging from neural and cognitive perspectives. Critical questions arise as to how such human-machine integration can be best achieved, given expected neurocognitive bottlenecks of brain plasticity. Here, we demonstrate that successful integration of motor augmentation can be achieved, with potential for flexible use, reduced cognitive reliance and increased sense of embodiment. Importantly, though, such successful human-robot integration may have consequences on some aspect of body representation and motor control which need to be considered and explored further.

## Methods

### Participants

36 healthy volunteers (23 females, mean age = 23.1 ± 3.89, all right handed) were recruited from internet-based advertisements and randomly assigned to either augmentation (n = 24, 14 females, mean age = 22.9 ± 4.12) or control (n = 12, 9 females, mean age = 23.5 ± 3.55) group. All participants were right-handed, between the ages of 18-35, did not have any known motor disorders and reported no counterindications for magnetic resonance imaging (MRI). Professional musicians were excluded from the study. Ethical approval was granted by the UCL Research Ethics Committee (REC: 12921/001). All participants gave their written informed consent before participating in the study.

Due to scheduling conflicts, 1 control participant and 3 augmentation participants dropped out of the study. Additionally, due to technical problems during data collection, 1 augmentation participant was discarded from the study.

### Experimental design

To assess the effects of hand augmentation on body representation, we implemented a longitudinal experimental design (see Figure 1C), involving 8 experimental sessions conducted across 7-9 days. All participants undertook (i) a 1-hour familiarisation session, introducing the equipment and the behavioural tasks; (ii), a 4-hour baseline (pre-test) session consisting of behavioural testing and an MRI scan; (iii) 5 2-hour training sessions conducted over the 5 subsequent days (1 session per day); (iv) a final 4-hour post-test session corresponding to the baseline session. Additionally, 12 of the participants from the augmentation group also undertook a secondary follow-up MRI session conducted 7-10 days after the end of training. Due to scheduling issues, 1 augmentation participant and 1 control participant completed only 4/5 training sessions.

All study participants were asked to wear an extra robotic thumb (the Third Thumb; Figure 1A) on their right-hand throughout the day. Participants were instructed to wear the Thumb during the in-lab training sessions and to continue wearing it outside of the lab for at least 4 hours per day. The augmentation group had full motor control over the Thumb and needed to actively use it to complete the training tasks. They were also encouraged to use it as much as possible outside of the lab for a free-style environment exploration (‘in the wild’). The control group wore a static (not-moving) version of the Thumb and completed the training tasks without being able to control it. Due to initial equipment issues, the first 2 control participants did not wear the Thumb during training.

### The Third Thumb

The Third Thumb is a 3D printed robotic thumb designed by Dani Clode (*17*) to extend the abilities of the biological hand by increasing the natural repertoire of finger synergies. The Thumb is worn over the ulnar side of the right palm, opposite to the user’s natural thumb (Figure 1A). It is actuated by two motors, allowing the control over 2 independent degrees of freedom - flexion/extension and adduction/abduction. The motors are mounted on a wrist strap (Figure 1B-1) and powered by an external battery pack worn around the arm (Figure 1B-2). The movement of the Thumb is controlled with pressure sensors taped underneath the big toes of the user’s feet. The pressure sensors are powered by the external batteries strapped around the ankles (Figure 1B-3). A wireless communication protocol is used to send the signal from the pressure sensors to the motors actuating the Thumb. Pressure exerted with the right toe controls flexion of the Thumb while the pressure exerted with the left toe controls the abduction. The extent of Thumb movement is proportional to the pressure applied.

### Usage measures ‘in the wild’

To monitor Thumb usage outside the lab, self-reported wear time and Thumb usage examples were collected daily from all wearers/users. Daily reports were averaged across days and an independent samples t-test was used to test for differences in wear-time between the augmented and control group. In addition, both pressure sensors were equipped with a SD-card data logger. While the Thumb was on, both sensors were logging the corresponding motor’s position and the associated timestamp to the SD cards. If the participant turned the Thumb off during the day, the recording was paused and resumed after the motor was restarted. Those recordings were used to further quantify the number of hours participants spent using the Thumb per day. Use time was defined as the time spent wearing the extra Thumb with the motors of the Thumb switched on, while movement time was defined as the time spent actively exerting pressure with the big toes while the Thumb was switched on. Due to initial equipment issues, the sensors’ data from first 3 augmentation participants were not recorded.

### Training protocol

During the training sessions, participants were asked to complete a set of reaching and grasping tasks. These tasks were designed to be purposefully challenging when performed with only one hand. The task execution was restricted to the augmented (right) hand. The augmentation group was instructed to use the extra Thumb to complete the training tasks. The control group, wearing the static version of the Thumb was instructed to complete the training tasks using only their natural fingers, i.e. without using the Thumb. For all of the tasks, participants were seated at a desk facing the camera recording their hand movements. Each task was conducted for 10-15 minutes and repeated on 2-4 separate training days, with the exception of the hand robot coordination task (see below), which was performed during each of the training sessions.

#### Collaboration tasks

In the collaboration tasks, participants had to use the extra Thumb in collaboration with another finger to pick up multiple objects. The collaboration tasks included building Jenga tower, sorting Duplo blocks, sorting shapes and manipulating multiple balls.

##### Building a Jenga tower

A mixed jumble of wooden Jenga blocks was placed in a shallow box. Augmentation participants were instructed to pick up 2 Jenga blocks at a time, using the Thumb to hold or support one of the blocks, and build a 2×2 Jenga tower. Control participants were instructed to use their biological fingers to hold and support both of the picked up Jenga blocks. The experimenter recorded the number of 2-block floors built in 1 minute.

##### Sorting Duplo blocks

Participants were presented with a set of colourful Duplo blocks and four colour-coded boxes. The goal of the task was to sort all the Duplo blocks into the matching colour boxes. Augmentation participants were instructed to pick up four blocks at a time (one of each colour), with one block being held or supported with the extra Thumb. Control participants were instructed to use their biological fingers to hold and support all of the Duplo blocks. The experimenter recorded the time taken to complete the task

##### Sorting shapes

Participants were given a wooden box with differently shaped holes and a set of wooden blocks matching the shapes. Picking up two blocks at a time, one with their biological fingers and one with the extra Thumb, augmentation participants had to sort the blocks into the corresponding holes. Control participants were instructed to used their biological fingers to hold and support both of the blocks. The experimenter recorded the time taken to complete the task.

##### Manipulating multiple balls

3 small foam balls were placed in shallow boxes in front of the participants. All participants were asked to pick up all 3 balls with their right hand, starting from the ball in the rightmost box, and to place them down in a different configuration. Augmentation participants were instructed to use the extra Thumb to pick up and hold one of the balls. Control participants were instructed to pick up and hold all of the balls with their biological fingers. The experimenter recorded the number of shuffles performed in 1 minute.

#### Shared supervision tasks

In the shared supervision tasks, participants had to use the extra Thumb to extend the natural grip of the hand and to free up the use of their biological fingers. Shared supervision tasks included picking up multiple wine glasses, plugging cables and stirring cups.

##### Wine glasses

5 plastic wine glasses were placed upside-down in front of the participants. All participants were instructed to pick up all 5 glasses with their right hand and to place them upright in a row, on the marked positions. Augmentation participants were instructed to use the extra Thumb to hold one of the glasses. Control participants were instructed to only use their biological fingers. The experimenter recorded the time taken to place all 5 of the glasses on the correct positions, the number of glasses that were dropped and the accuracy of the placement.

##### Plugging cables

Participants were given 4 long USB cables and a USB hub with 4 separate ports. All participants were instructed to pick up the USB hub and plug in all 4 USB cables, while holding the hub in the air. Augmentation participants were instructed to use the extra Thumb to complete the task. Control participants were instructed to only use their biological fingers. The experimenter recorded the time needed to plug in all of the USB cables.

##### Stirring cups

3 small marbles were placed in 3 plastic cups. All participants were asked to pick up one cup at a time and scoop out the marble with a plastic spoon, whilst holding the cup in the air. Augmentation participants were instructed to hold the cup using the Thumb. Control participants were instructed to only use their biological fingers. After the first day of training, the cups were filled with the Styrofoam balls to increase the difficulty of the task. The experimenter recorded the time taken to scoop out each of the marbles.

#### Robotic Thumb individuation task

##### Stacking tape rolls

In the individuation task participants had to work on the fine motor control of the Thumb, while having their hand fully occupied with a task-irrelevant object. Specifically, participants were given 6 tape rolls, a foam ball and a wooden pole fixed to the desk. While holding the ball with their biological fingers, augmentation participants had to use the extra Thumb to pick up the tape rolls and place them on the wooden pole. Control participants were instructed to use their biological thumb to pick up the tapes while holding the ball with the remaining digits. The experimenter recorded the time taken to place all the tapes on the wooden pole and the number of dropped tapes per trial.

#### Hand-Thumb coordination (Thumb opposition)

To monitor daily changes in hand-Thumb coordination, we used a finger opposition task (previously used to monitor hand function in healthy and clinical cases, as well as robotic finger operation, (*4, 12, 13*)). This task was conducted at the start of each training session. Participants were seated in front of a computer screen that displayed task stimuli.

Augmentation participants were instructed to move the Thumb to touch the tip of a randomly specified finger on the augmented (right) hand. Control participants performed a modified version of this task in which they were instructed to use their biological thumb to oppose the remaining digits of the right hand. All participants were instructed to attempt to make as many successful hits as possible within a 1 minute block. Participants completed a total of 10 blocks. A MATLAB script was used to randomly select a target finger (thumb, index, middle, ring or little) and to display the finger name on the computer screen in front of the participant. The experimenter manually advanced the program to the next stimulus when the participant successfully touched the tip of the target finger with the extra Thumb or with biological thumb (hit); or when a wrong finger has been touched (miss).

To quantify the improvement of the augmentation group on each of the training tasks, the outcome measure of each training task was averaged for each participant and each training session. As different participants had slightly different training regimes, in terms of distribution of tasks across the days, we sorted the average scores based on the order of task repetition (i.e. 1^st^, 2^nd^, 3^rd^ time the task was repeated regardless of which days it was repeated on). These data were then analysed using a repeated measures ANOVA in SPSS. One of the shared supervision tasks (plugging cables) was not analysed due to inconstancies in task execution.

#### Numerical cognition

To assess the cognitive load related to Thumb use, a numerical cognition task was performed twice, on both the first and the last training sessions (*40–42*). Participants were asked to perform a cooperation task, building a Jenga tower (described above), while simultaneously presented with a set of low and high pitch auditory tones played from a laptop. The tones were presented every 1-6s in a randomised order, for a total duration of 1 minute per block. Starting with a number 10, participants were instructed to add 1 to the current number after hearing a high tone, and subtract 1 from the current number after hearing a low tone After each mathematical operation, participants were instructed to verbally respond with the resulting number. Participants performed 5 blocks of the numerical cognition task during each session. Numerical cognition blocks were always preceded and followed with 5 blocks of normal (baseline) building a Jenga tower task (5 baseline blocks, 5 numerical cognition blocks, 5 baseline blocks). Note that the first 3 participants did not complete the numerical cognition task.

For each participant, the average number of Jenga floors built was calculated from all the numerical cognition blocks in which the correct mathematical operations were performed. Trials in which a wrong number was given were discarded (on average 15-19% of trials were discarded per participant). To determine whether the extra cognitive load caused by the numerical cognition task had any impact on participants’ motor performance when using the extra Thumb, the average score from the numerical cognition task was compared to the baseline score. This was done separately for the first and the last day of training. The baseline score was calculated as the average number of Jenga floors built across the two blocks (10 trials) that proceeded and followed the numerical cognition task.

### Tracking hand movements

To assess the changes in finger coordination across all training tasks, we tracked the kinematics of the augmented (right) hand using flex sensors embedded in a dataglove (CyberGlove, Virtual Technologies, Palo Alto, CA, USA). Finger kinematics have been previously shown to reflect the brain organisation better than EMG-derived measures (*9*). Here, the sensors of the dataglove were associated with 19 degrees of freedom of the hand and included the metacarpal-phalangeal (MCP), proximal interphalangeal (PIP) and distal interphalangeal (DIP) joint angles for the four fingers (index, middle, ring and little finger), the carpometacarpal, (CMC) metacarpal-phalangeal (MCP) and interphalangeal joint (IP) angles for the biological thumb, the three relative abduction angles between the four fingers and the abduction angle between the biological thumb and the palm of the hand. Sensors were sampled continuously at 100 Hz using Shadow Robot’s (https://www.shadowrobot.com) CyberGlove interface for the Robot Operating System (ROS, https://www.ros.org).

Participants wore the CyberGlove underneath the extra Thumb throughout all of the training sessions. Kinematics associated with each of the training tasks performed during a given session were recorded onto a separate file. Due to initial equipment issues, the first 4 augmentation participants did not wear the CyberGlove during training.

#### Cyberglove calibration

The CyberGlove was calibrated for each participant at the beginning of each training session, using a min-max pose calibration procedure provided with the ROS CyberGlove package. During the calibration, participants were presented with a set of carefully selected hand poses and given a few seconds to shape their hand accordingly. For each hand pose, the sensor values were sampled and averaged over one second of recording. Averaged sensor values were then saved alongside the actual joint angles, determined based on the hand configuration. Once all hand poses have been recorded, a linear regression was used to calculate the mapping from the sensor-values to the joint angles.

#### Hand kinematics analysis

We focused the hand kinematics analysis on the data recorded during the first and last days of training. Recorded data from the 19 sensors were calibrated using the established mapping from the sensor-values to the joint angles. The joint angles were smoothed using a 3^rd^ order Savitzky-Golay filter, with a window length of 151 samples. Angular velocities were then calculated from the first difference of the filtered joint angle data divided by the time step. Since most of the finger movements employed during the training tasks were executed using the PIP joints, to simplify the analysis (*9*), only data from these five joints were analysed. Due to acquisition errors, for 2 augmentation participants and 2 control participants, the data recorded during the first day of training was unavailable. Therefore, for these participants, the data from the second day of training was used instead. Similarly, when the data from the last day of training was unavailable (3 augmentation participants, 2 control participants), the data from the penultimate day of training was used instead. Due to calibration issues, the hand kinematics data of 1 augmentation and 2 control participants were discarded from subsequent analysis.

To quantify the complexity of the hand movements across both groups, we first conducted a Principal Components Analysis (PCA) of the angular velocities of the PIP joints. Prior to PCA, the angular velocities were z-normalised (*43*) note however that similar results were obtained with not normalised data (*44*). The extracted principal components (PCs) were ordered according to the amount of variance explained by each component. Consistent with the literature (*44, 45*), we found that the first PC accounted for more than 40% of total variance and reflected a coordinated movement of all the fingers (Figure 2A and Supplementary Figure S3 for all the PCs). To quantify the dimensionality of the hand movements, for each participant and day, we recorded the number of PCs needed to explain 80% of total variance (*46*). These were then compared across groups in a repeated-measures ANOVA with time (day 1, day 5) as a within-subjects factor and group (augmented, control) as between-subject factor. Next, to quantify the contribution of the all-digit movements to the complexity of the hand kinematics, we compared the amount of variance explained by the first PC across both groups using the same repeated-measures ANOVA design.

Next, to interrogate more detailed changes to the finger cooperation pattern caused by the hand augmentation, we looked at the degree of coupling between digit pairs, adapting the methods used in (*44*). We used linear regression to fit the angular velocity data of a given digit as a function of the angular velocity of each of the other digits individually. This yielded a single determination coefficient (R^2^) for each digit pair, expressing the proportion of total variance of each digit’s angular velocity that could be explained by a linear reconstruction, based on its paired regression with each of the other four digits. Qualitative comparison between the results obtained in the control group (who did not use the extra Thumb during the training) and outcomes of previous hand kinematics research conducted during free movement (*44*), confirmed that the conducted analysis resulted in a typical finger coordination profile.

To assess the effect of Thumb use on the finger coordination profile across groups, we performed a linear mixed model analysis (LMM) with fixed factors of time, group (augmentation vs controls) and digit pair, a random effect of participant and a random participant-specific slope of time. The LMM was evaluated in R (version 3.5.2) under restricted maximum likelihood (REML) conditions with Satterthwaite adjustment for the degrees of freedom.

### Pre-post testing protocol

To assess the neural correlates of hand augmentation we used a set of pre- to post-training comparison measures, consisting of both behavioural and neuroimaging tasks. To characterise the emerging representation of the extra Thumb, we probed the proprioception and motor control of the Thumb using a sequential variation of the hand-Thumb coordination task. We also assessed perceived (phenomenological) embodiment of the Thumb using questionnaires. To interrogate changes to the natural hand representation, we measured biological finger co-dependencies using finger kinematics and force-enslavement task. Changes to body image were probed using tactile distance and hand laterality judgement tasks. Finally, fMRI was used to track the hand representation in the sensorimotor cortex of the brain and to interrogate changes to the relationship between the hand and feet representations. With the exception of the hand-Thumb coordination task, participants were not wearing the Thumb during testing.

#### Hand-Thumb coordination (sequential)

To probe changes to implicit motor control of the Thumb, a sequential variation of the hand-Thumb coordination (finger to Thumb opposition) task has been used. In this task, participants sequentially opposed the Thumb to the tip of each of the five fingers of their augmented hand, starting with the little finger. Participants were instructed to repeat this movement cycle as many times as possible within a 1-minute block, while maintaining high accuracy. The task consisted of 5 blocks. To assess the proprioception of the Thumb, participants were further asked to perform 5 blocks of the same task while blindfolded. The experimenter recorded the number of successful hits per block. For each participant, an average score (number of hits) was calculated separately for each session (pre, post) and vision condition (with vision, blindfolded). Due to a data acquisition mistake, 1 control participant was not included in the analysis.

#### Embodiment questionnaires

To assess changes in the phenomenological embodiment of the Thumb, participants were asked to complete an embodiment questionnaires before the first training session and again after the last training session. Due to data collection issues, 5 augmentation participants and 1 control participant only completed the post-training embodiment questionnaire. Participants were asked to rate their agreement with 12 statements (based on (*19*)) on a 7-point Likert-type scale ranging from −3 (strongly disagree) to +3 (strongly agree). Statements were clustered into 4 main categories, probing different aspects of embodiment, namely: body ownership, agency, body image and somatosensation.

**Table 1.**
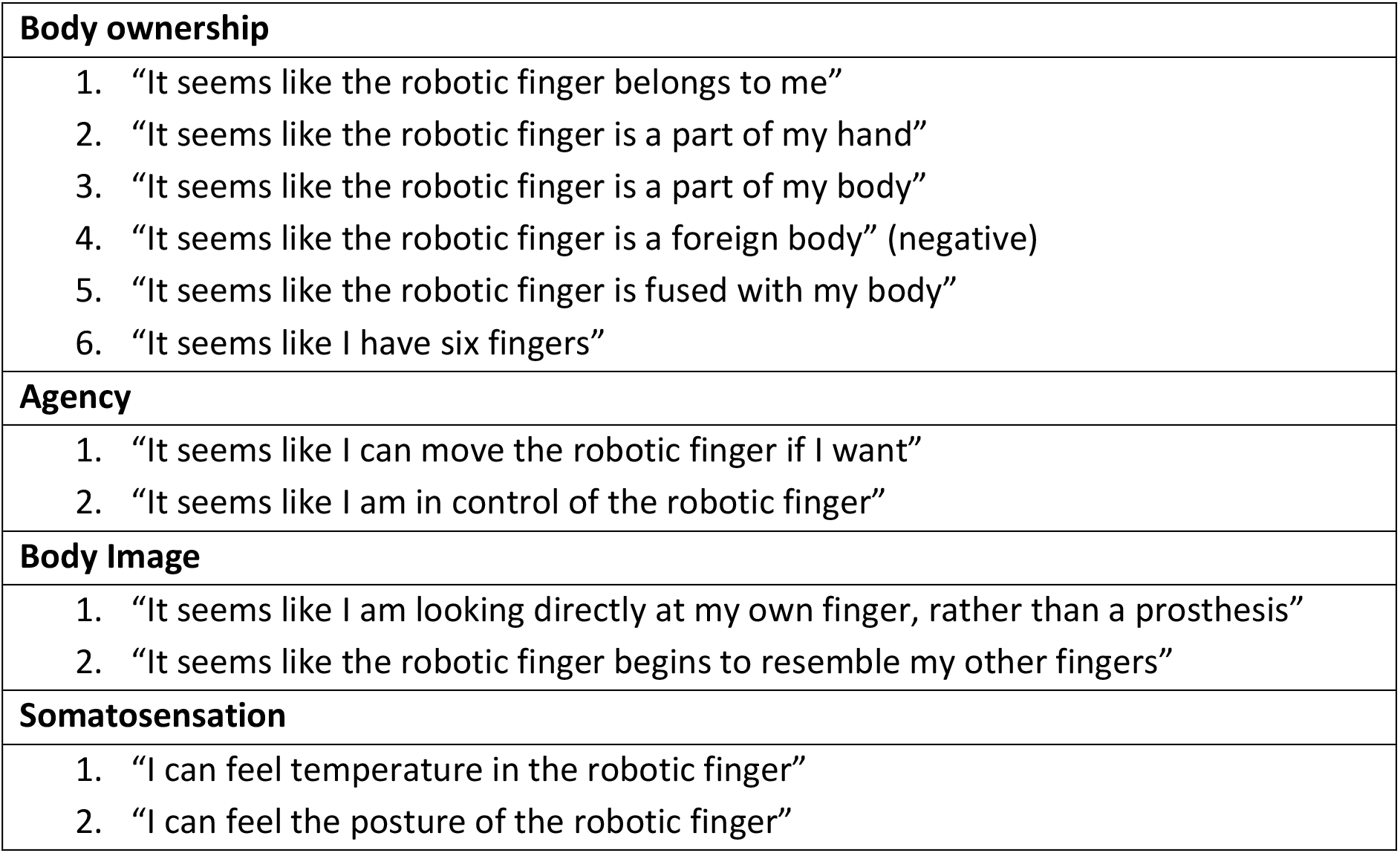
Embodiment questions divided into 4 separate embodiment categories

|  |
| --- |
| **Body ownership** |
| 1. "It seems like the robotic finger belongs to me"<br>2. "It seems like the robotic finger is a part of my hand"<br>3. "It seems like the robotic finger is a part of my body"<br>4. "It seems like the robotic finger is a foreign body" (negative)<br>5. "It seems like the robotic finger is fused with my body"<br>6. "It seems like I have six fingers" |
| **Agency** |
| 1. "It seems like I can move the robotic finger if I want"<br>2. "It seems like I am in control of the robotic finger" |
| **Body Image** |
| 1. "It seems like I am looking directly at my own finger, rather than a prosthesis"<br>2. "It seems like the robotic finger begins to resemble my other fingers" |
| **Somatosensation** |
| 1. "I can feel temperature in the robotic finger"<br>2. "I can feel the posture of the robotic finger" |

For each participant, questionnaire scores were averaged within each embodiment category. Note that in the body ownership category, the opposite (negative) value of the ‘foreign body’ statement has been used while computing the average. 1 augmentation participant was discarded from this analysis, as their averaged agency score was classified as a statistical outlier (different from the mean score by more than 3 standard deviations).

#### Force enslavement

To estimate the degree of co-dependency across the biological fingers, a custom-built keyboard (previously used in (*9*)) was used to measure isometric finger forces generated during individuated finger presses.

During the pre and post testing sessions, participants performed individuated force presses with instructed fingers, while forces produced with all of the fingers were recorded. Participants were instructed to always keep all of their fingers on the keys of the keyboard. A visual representation of the force exerted with all 5 fingers was presented on a screen (Figure 3C). Following the methods described in (*9*), the experiment began by estimating the strength of each finger, by measuring two repetitions of the maximum voluntary force (MVF) of each digit. All subsequent trials required the production of isometric fingertip forces at a 75% of the maximum voluntary force for the instructed digit (*9*). At the start of every trial, a force target-zone (target-force ± 25%) was highlighted in green in the visual display. This cued participants to make a short force press with the instructed finger in order to match and maintain the target-force for 1s while keeping the uninstructed fingers as stable as possible. The trial was stopped if force of the instructed digit did not reach the target-zone within the 2s following the stimulus onset. Single trials, presented in a randomised order were grouped in blocks, with each block consisting of 40 trials (8 trials per finger). Participants performed 4 force-enslavement blocks during each session. The data of two augmentation participants were excluded from the subsequent analysis, as those participants produced extraordinarily low MVF forces (below 2N).

For each participant, the recorded force data was first filtered with a 3^rd^ order Savitzky-Golay filter with a window length of 51 samples. The data was then separated into individual trials (finger presses). Trials in in which the force produced with the instructed finger did not reach the target force were discarded from the analysis. Within each trial, a linear regression was used to fit the force generated by the instructed digit as a function of the force generated by each of the other digits. The resulting determination coefficients (R2) were averaged across trials to yield a 5×5 matrix force enslavement matrix, expressing the involuntary force changes across non-instructed fingers during the presses of the instructed finger.

To estimate gross changes to finger co-dependency, we performed a linear mixed model analysis (LMM) with fixed factors of time, group (augmentation vs controls) and digit pair, a random effect of participant and a random participant-specific slope of time. The LMM was evaluated using the same parameters as the ones used for finger co-use modelling (see Hand kinematics analysis). To test the assumption that having an extra robotic Thumb would impact the independence of the biological thumb, a similar linear mixed model was created using only the force enslavement caused by the thumb (4 values, one for each enslaved finger).

#### Tactile distance perception

To examine whether Thumb usage would lead to incorporating it into the body representation, we tested participants’ ability to discriminate between tactile distances applied over their wrist and forearm. The design was inspired by a previous study (*47*) showing greater biases in spatial tactile perception when tested across a joint. Prior to the experiment, the experimenter marked the midpoint of the participant’s forearm, as well as the basepoint of the extra Thumb with a pen.

During the experiment, participants were seated in an armchair, with their right elbow rested on an elevated foam padding with the forearm at full flexion and their left hand placed on a mouse connected to the experimental computer. The tactile stimuli (hereafter distances) comprised of custom-made callipers with acrylic pins fixed at distances of 50, 60 and 70 mm. In each trial, two distances were presented sequentially – one over the marked basepoint of the Thumb, one over the midpoint of the ventral side of the forearm (both in the same orientation). The experimenter presented the distances manually ensuring that the two points of each calliper touched the skin simultaneously. Participants were instructed to indicate which of the distances they perceived as larger by pressing the left (distance over the Thumb’s base perceived as larger) or right (distance over the forearm perceived as larger) mouse button with their left hand. The task consisted of 3 blocks. Each block included 5 presentations of each of the following ratios of distances in a randomised order: 50/70mm, 50/60mm, 60/70mm, 60/60mm, 70/60mm, 60/50mm, and 70/50mm).

Following the methods described in (*47*), we measured the proportion of responses in which the stimuli applied over the basepoint of the Thumb was judged as larger, as a function of the ratio of the length of the stimuli. Cumulative Gaussian curves were fit to the data using Matlab (v. 2017b). Point of subjective equality (PSE) was calculated separately for each participant and session, as the ratio of stimuli at which the psychometric function crossed the 50%. Additionally, the interquartile range (IQR) – that is the difference between the points on the x-axis where the psychometric function crosses 25% and 75% - was calculated as a measure of the precision of participant’s judgements. One augmentation participant and two control participants were excluded from further analysis due to poor goodness of fit coefficient (R^2^<0.15) for the psychometric function in the pre-session. The remaining R^2^ scores, averaged across participants, showed an excellent fit to the data (R^2^=0.88±0.15

#### Hand Laterality Judgement

To examine whether Thumb usage leads to changes in body image, participants were presented with drawings of hand outlines, adapted from (*48*) and asked to decide whether the hand drawings corresponded to a right or a left hand. Participants were instructed to respond verbally by indicating the hand laterality (left or right) of each presented image as fast as possible, while maintaining high accuracy. The stimuli included drawings of left and right hands, presented in four different postures (dorsal view, palm view, side from thumb view, and palm from wrist view) and at 7 different angles (upright 0°, 30°, 90°, 150°, 210°, 270°, and 330° in a clockwise direction). Participants completed four experimental blocks, each including all of the 56 hand images. Hand images were presented in a random order using Psychopy software. Participants were seated comfortably in front of a laptop computer with their hands obstructed by a black cape. Each hand drawing was preceded by 1 second presentation of a fixation cross and disappeared either after a verbal response was provided or after 10 seconds of no response. Time from the start of the image display to voice onset was recorded as the participants’ reaction time (RT). Audio files with participant’s responses were recorded for off-line accuracy analysis. Due to equipment malfunction, 2 augmentation participants and 1 control participant did not complete the hand laterality judgement task.

All audio recordings and the appropriate classification of reaction-times were verified offline by a naive experimenter. Trials with noisy recordings were excluded from further analysis. Accuracy was computed as the proportion of correct response of all valid trials. Only trials with correct responses were included in the RT analysis. RTs were logarithmically transformed in order to correct for the skewed RT distribution and to satisfy the conditions for parametric statistical testing. Transformed RTs deviating more than 3 standard deviations from the participants’ means (separately for each session) were discarded.

### Statistical analysis

All statistical analysis was performed using IBM SPSS Statistics for Macintosh (Version 24), R (for linear mixed models) and JASP (Version 0.11.1). Tests for normality were carried out using a Shapiro-Wilk test. Training data that were not normally distributed were log-transformed prior to further statistical analysis. With the exception of hand kinematics and force enslavement datasets, that were analysed using linear mixed models (LMM), all the within group comparisons were carried out using paired t-tests or repeated measures ANOVAs (training tasks data). Between group comparisons were carried out using ANCOVAs with group (augmentation, controls) as a fixed effect and the pre-score used as a covariate (*49*). All non-significant results were further examined using corresponding Bayesian tests.

### Scanning procedures

Both pre- and post- neuroimaging sessions were comprised of the following functional scans: (i) a resting state scan, (ii) a motor localiser scan and (iii) four finger-mapping scans. Additionally, a structural scan and field maps were obtained during each scanning session.

#### Resting state scan

Participants were instructed to let their mind wander while keeping their eyes loosely focused on a fixation cross for the duration of the scan (5 min).

#### Motor localiser scan

Participants were instructed to move the right/left hand (all fingers flexion and extension), right/left foot (unilateral toe movements), or lips (blowing kisses) as paced by visual cues projected into the scanner bore. Each condition was repeated four times in a semi-counterbalanced protocol alternating 12s of movement with 12s of rest. Participants were trained before the scan on the degree and form of the movements. To confirm that appropriate movements were made at the instructed times, task performance was visually monitored online. Due to data acquisition issue, motor localiser data of 1 augmentation participant was discarded from the analysis.

#### Finger-mapping scans

Participants were instructed to perform visually cued movements of individual digits of either hand, bilateral toe movements and lips movements. The different movement conditions, as well as rest periods were presented in 9s blocks. The individual digit movements were performed in the form of button presses on MRI-compatible button-boxes (4 buttons per box) secured on the participant’s thighs. The movements of either of the (biological) thumbs were performed by tapping them against the wall of the button box. Instructions were delivered via a visual display projected into the scanner bore. Ten vertical bars, representing the fingers flashed individually in green at a frequency of 1 Hz, instructing movements of a specific digit at that rate. Toe and lips movements were cued by flashing the words “Feet” or “Lips” at the same rate of 1 Hz. Each condition was repeated 4 times within each run in a semi-counterbalanced order. Participants performed 4 runs of this task. Due to timing issues 3 augmentation participants and 1 control participants completed only 3 runs of the finger-mapping task. Additionally, due to a data acquisition issue, the finger-mapping data of 1 control participant was discarded.

### MRI data acquisition

MRI images were acquired using a 3T Prisma MRI scanner (Siemens, Erlangen, Germany) with a 32-channel head coil. Functional images were collected using a multiband T2*-weighted pulse sequence with a between-slice acceleration factor of 4 and no in-slice acceleration. This provided the opportunity to acquire data with high spatial (2 mm isotropic) and temporal (TR: 1450 ms) resolution, covering the entire brain. The following acquisition parameters were used: TE: 35ms; flip angle: 70°, 72 transversal slices. Field maps were acquired for field unwarping. A T1-weighted sequence (MPRAGE) was used to acquire an anatomical image (TR: 2530 ms, TE: 3.34 ms, flip angle: 7°, spatial resolution: 1 mm isotropic).

### MRI analysis

MRI analysis was implemented using tools from FSL (*50, 51*) and Connectome Workbench (humanconnectome.org) software, in combination with Matlab scripts (version R2016a), both developed in-house (including FSL-compatible RSA toolbox (*52*)) and as part of the RSA Toolbox (*21*). Cortical surface reconstructions were produced using FreeSurfer ((*53, 54*), freesurfer.net).

### fMRI pre-processing

Functional data was first pre-pre-processed using FSL-FEAT (version 6.00). Pre-processing included motion correction using MCFLIRT (*55*), brain extraction using BET (*56*), temporal high pass filtering, with a cut off of 150s for the finger-mapping scans and 100s for resting-state and motor localiser scans, and spatial smoothing using a Gaussian kernel with a FWHM of 3mm for the finger-mapping and 5mm for resting-state and motor localiser scans.

To make sure that the scans from the two scanning sessions were well aligned, for each participant we calculated a midspace between their pre- and post- scans, i.e. the average space in which the images are minimally reoriented. Each scan was then aligned to this prepost midspace using FMRIB’s Linear Image Registration Tool (FLIRT, (*55, 57*)). This registration was performed separately for the structural, motor localiser, resting state and finger mapping scans, resulting in 4 separate midspaces per participant. All the within-subject analyses were done in the corresponding functional midspace. To allow for the co-registration of the functional data and the anatomical ROIs (see below) the functional midspaces were then aligned to each participant’s structural midspace using FLIRT, optimised using Boundary-Based Registration (BBR, (*58*)). Finally, when interrogating finger-specific activations on the group-level, the individual structural midspaces were transformed into MNI space using FMRIB’s Nonlinear Image Registration Tool (FNIRT, (*57*))

### Low-level task-based analysis

For task-based datasets, voxel-wise General Linear Model (GLM) analysis was carried out using FEAT, to identify activity patterns related to the movement of each digit/body part. The design was convolved with a double-gamma hemodynamic response function (HRF) and its temporal derivative. The six motion parameters were included as regressors of no interest. In case of large movement between volumes (>1 mm) additional regressors of no interest were included in the GLM to account for each of these instances individually.

For the finger-mapping scans, 12 contrasts were set up: each digit versus rest, feet against rest and lips against rest. The estimates from the four finger mapping scans were then averaged voxel-wise using fixed effects model with a cluster forming z-threshold of 3.1 and family-wise error corrected cluster significance threshold of p<0.05, creating 12 main activity patterns for each session and participant.

For the motor localiser scans, 4 main contrasts were set up: right/left hand against lips, rand right/left foot against lips. The activity patterns associated with those 6 contrasts were then used to define functional regions of interest (ROIs, see below).

### Regions of Interest (ROI) definition

Changes to representational structure of the hand were studied using anatomical ROIs, as previously practiced in related studies (*13, 59, 60*). Structural T1 images, registered to the structural midspace, were used to reconstruct the pial and white-grey matter surfaces using Freesurfer. Surface co-registration across hemispheres and participants was done using spherical alignment. Individual surfaces were nonlinearly fitted to a template surface, first in terms of the sulcal depth map, and then in terms of the local curvature, resulting in an overlap of the fundus of the central sulcus across participants (*61*). The anatomical sensorimotor ROI, used for the multivariate analysis were defined on the group surface using probabilistic cytotectonic maps aligned to the average surface (*62*). These ROI was then projected into the individual brains via the reconstructed individual anatomical surfaces. Since we were primarily interested in the motor representation of the hand, we have focused our anatomical ROI on M1, selecting all surface nodes with the highest probability for BA4 spanning a 2cm strip medial/lateral to the anatomical hand knob (*13, 63*). However, we note that, given the probabilistic nature of these masks, the dissociation between S1 and M1 is only an estimate, and thus our ROI should be treated as a sensorimotor one.

For our univariate analyses, we also defined functional ROIs based on the sensorimotor representations of the left/right hand and feet for each participant. Unlike the cross-validated RSA analysis, these analyses require more spatially-restricted ROIs. We therefore used condition-specific contrasts from the motor localiser scans (as previously practiced in e.g. (*12, 64–66*) To this extend, the relevant (right/left hand vs lips, right/left foot vs lips) low-level contrasts were first averaged across both (pre and post) scans. To create left/right hand ROIs, for each participant, the 200 most active voxels were selected from the averaged left/right hand vs lips contrast. For the individual feet ROIs, a similar procedure was employed, selecting the 100 most active voxels from the averaged left/right foot vs lips contrast. Voxel selection was restricted to the primary somatosensory (S1) and motor (M1) cortices as derived from the maximum probabilistic maps (thresholded at 25%) of the Juelich Histological Atlas (*67*). Voxels from both feet ROIs were combined into a single region of interest used in the subsequent analyses.

### Resting state

To account for non-neuronal noise that might bias functional connectivity analyses (*68, 69*), we regressed out the six motion parameters, as well as the BOLD time series of white-matter and cerebrospinal fluid (CSF) from the pre-processed resting state data. For this purpose, the T1-weighted structural scans, registered to anatomical midspace, were segmented into grey matter, white matter, and CSF, using FSL FAST (*70*). To avoid the inclusion of grey matter voxels in the nuisance masks, the resulting masks included only voxels identified as white matter/CSF with probability of 1, and were eroded by one voxel in each direction. For both the white matter and CSF maps, the first five eigenvectors were then calculated using the unsmoothed resting state time series (*68, 69*). Together with the motion parameters, these 16 regressors of no interest were regressed out from the pre-processed resting state time series. The resulting time series (residuals) were used in all the subsequent resting state analyses.

The level of resting-state coupling (functional connectivity) between the augmented (right) hand ROI and the feet ROIs was examined by correlating the time-course of the augmented hand ROI with the time-course of the feet ROI, while partialling out the time-course of the left-hand ROI. The resulting sets of partial correlations were Fisher z-trasformed, and group-level statistical comparisons were conducted using two-tailed paired t-tests.

### Multivariate representational structure (RSA)

We used RSA (*71*) to assess the multivariate relationships between the activity patterns generated across digits and sessions. The dissimilarity between activity patterns within the M1 anatomical hand ROI was measured for each digit pair using the cross-validated squared Mahalanobis distance (*21*). We calculated the distances using each possible pair of imaging runs within a single scanning session (pre, post) and then averaged the resulting distances across run pairs. Before estimating the dissimilarity for each pattern pair, the activity patterns were pre-whitened using the residuals from the GLM. Due to the cross-validation procedure, the expected value of the distance is zero (or below) if two patterns are not statistically different from each other, and larger than zero if the two representational patterns are different. The resulting 10 unique inter-digit representational distances were put together in a representational dissimilarity matrix (RDM).

To assess the effect of 5-day Thumb usage on the overall representation structure (dissimilarity), we performed a linear mixed model analysis (LMM) with fixed factors of time, hand and digit pair, a random effect of participant and a random participant-specific slope of time. The LMM was evaluated in R (version 3.5.2) under restricted maximum likelihood (REML) conditions with Satterthwaite adjustment for the degrees of freedom. We further assessed the typicality of the representational structure of the post scan RDM, by calculating the Spearman’s rho correlation between each participant’s RDM and the group average computed from the pre-scan data using a leave-one-out procedure. The typicality values were then z-normalised and the typicality of the representational structure of the right hand (postscan) was then compared to the typicality of the left hand’s representational structure (postscan) using paired t-test. Because the representational structure can be related to behavioural aspects of hand use and is highly invariant in healthy individuals (average correlation r = 0.9, (*9*)), this measure serves as a proxy for how ‘normal’ the hand representation is.

As an aid to visualising the RDMs, we also used classical multidimensional scaling (MDS). MDS projects the higher-dimensional RDM into a lower-dimensional space, while preserving the inter-digit dissimilarity values as well as possible. MDS was performed on data from individual participants and averaged after Procrustes alignment (without scaling) to remove arbitrary rotation induced by MDS. Note that MDS is presented for intuitive visualisation purposes only, and was not used for statistical analysis.

### Numerical modelling of inter-digit dissimilarity

To aid the interpretation of the neuroimaging findings, we have created a simple numerical simulation, modelling the potential effects of: (i) cortical magnification, (ii) inhibition and (iii) adding a new digit representation; onto the canonical hand representational structure.

We aspired to simulate activity patterns within the hand ROI by creating an abstract structure (comprised of 3000 “voxels”), divided into 5 equisized finger-selective regions, and simulating activity elicited by individuated movements of each of the fingers. During each trial (simulation run), the moving finger activated the “voxels” within the assigned finger-selective region (with inherent noise), while also partially activating the “voxels” assigned to the other 4 fingers. More specifically, within a trial, the activity of each “voxel” was randomly sampled from a Gaussian distribution, with mu=1 for trials involving the “voxel’s” preferred finger and ranging from 0.1408 to 0.3846 for trials involving a movement of a different finger. The values of mu were chosen based on the inter-finger relationship derived from the average representational structure (RDM) of the dominant hand of an independently acquired cohort of participants (*13*). The noise level (sigma=0.5) was constant across all trials and activations. The simulation was run 10,000 times, separately for each of the 5 fingers, resulting in 50,000 patterns of finger specific “activations”.

To model the cortical magnification phenomenon, we increased the mu values associated with all the non-moving fingers by 10% (activation modifier, see Table 2). This introduced the idea that increased individuation of the finger movements (based on Figure 2 above) results in increased representation of the fingers (*37*) Similarly, modelling the homeostatic inhibition, we decreased all the mu values by 10% (see Table 2). This interrogates the idea that any changes triggered by the changed finger co-use would be offset by tonic inhibition, that will impact the entire hand map (*36*). Finally, to investigate the theoretical effect of accounting for a new (6^th^) digit representation added into the hand representational structure, we decreased the number of “voxels” assigned to each of the 5 fingers, in order to create a new equisized digit-specific region. Since we didn’t have any a priori assumptions on the representational relationship between the extra Thumb and the rest of the fingers, this area was set to be activated equally in all trials, with mu set to an average activation value for the not-moving fingers (sigma=0.5, mu=0.2193).

**Table 2.** Parameters used for the numerical simulation of cortical magnification, homeostatic inhibition and addition of a new digit.

| **Parameters / simulation** | **Canonical** | **Cortical magnification** | **Homeostatic inhibition** | **New digit** |
| --- | --- | --- | --- | --- |
| Number of voxels per digit | 600 | 600 | 600 | 500 |
| Noise level (sigma) | 0.5 | 0.5 | 0.5 | 0.5 |
| Mean activation of the moving finger | 1 | 1 | 0.9 | 1 |
| Activation modifier for the non-moving fingers | 1 | 1.1 | 0.9 | 1 |

For each simulation run and each model, we computed the Euclidean distances between the activity patterns associated with each digit’s movement. We then averaged the distances across all 10 digit pairs to obtain an average dissimilarity measure. Finally, we compared the average distances computed for each of the models to the ones calculated for the canonical hand model, using independent samples t-tests.

## Acknowledgements

This work was supported by an ERC Starting Grant (715022 EmbodiedTech), awarded to TRM, who was further funded by a Wellcome Trust Senior Research Fellowship (215575/Z/19/Z) and by Sir Halley Stewart Charitable Trust (580). We thank Dominic Stirling, Samuel Cousins, Lydia Mardell, Maria Kromm and Mathew Kollamkulam for their help with data collection; Ekaterina Tupitsyna for developing the script for the kinematic data analysis; James Kilner for providing us access to the Cyberglove; Joern Diedrichsen for the custom-made force keyboards; Howard Bowman for introducing us to information theory measures; Gionata Salvietti for invaluable technical help during piloting; Silvestro Micera, Juan Alvaro Gallego and Daan Wesselink for helpful comments on the manuscript; Hunter Shone for proof-reading the manuscript; Domenico Prattichizzo for inspiring discussions about supernumerary fingers; and our participants for taking part in this study.

