## Supplementary Information for "Neurocognitive consequences of hand augmentation"

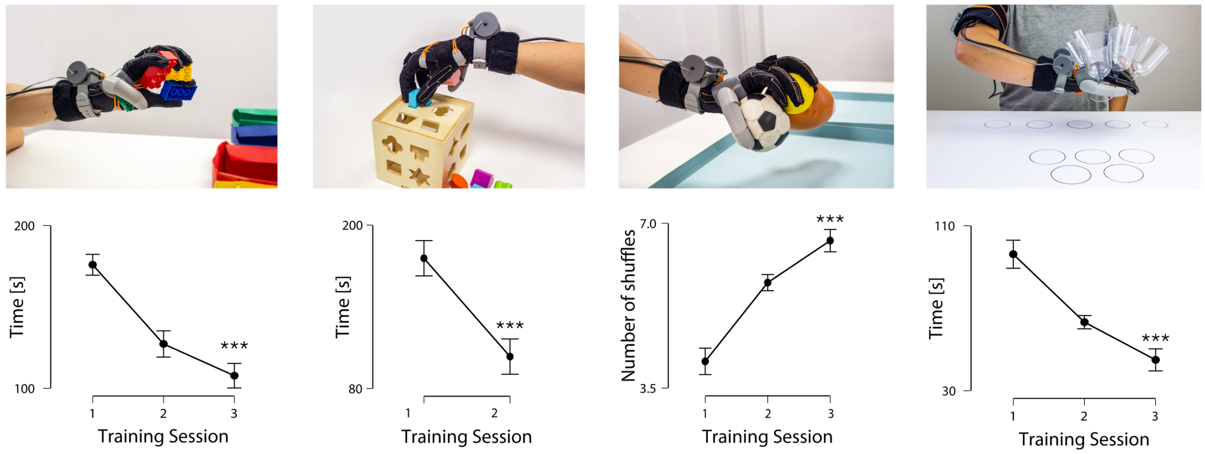


**Supplementary Figure S1**. Examples of the in-lab training tasks used for hand-Thumb collaboration (sorting Duplo blocks, sorting shapes, manipulating multiple balls) and shared supervision (wine glasses). Participants showed significant performance improvements across all training session. Asterisks denote significant main effect of time at *** p<0.001.

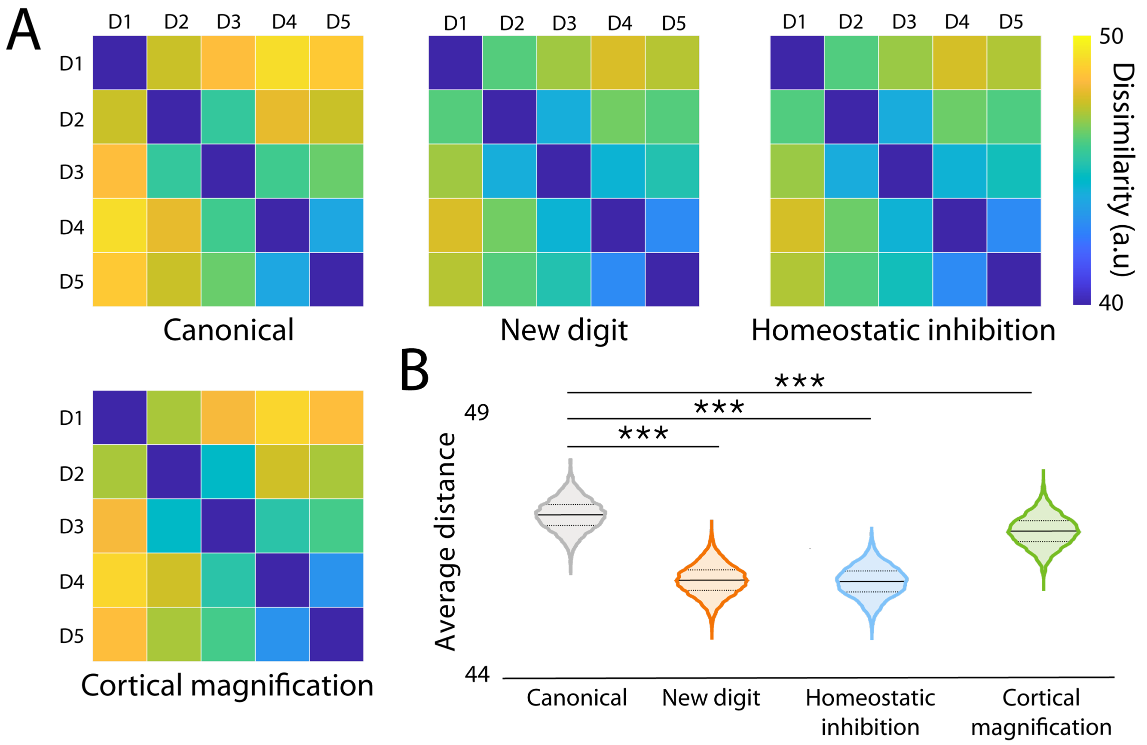


**Supplementary Figure S2**. To aid the interpretation of the neuroimaging findings, we have created a simple numerical simulation, modelling the effects of (i) cortical magnification, (ii) inhibition and (iii) adding a new digit representation on the canonical hand structure. (A) Mean dissimilarity matrices computed from 10000 simulations of each of the models. (B) Average distance (dissimilarity) is significantly decreased, as compared to the canonical hand representation, by homeostatic inhibition, adding a representation of a new digit and cortical magnification. Solid lines represent the mean of 10000 simulations, dashed lines denote the 1$^{st}$ and 3$^{rd}$ quartile of the data. Asterisks denote significant effects at *** p<0.001.

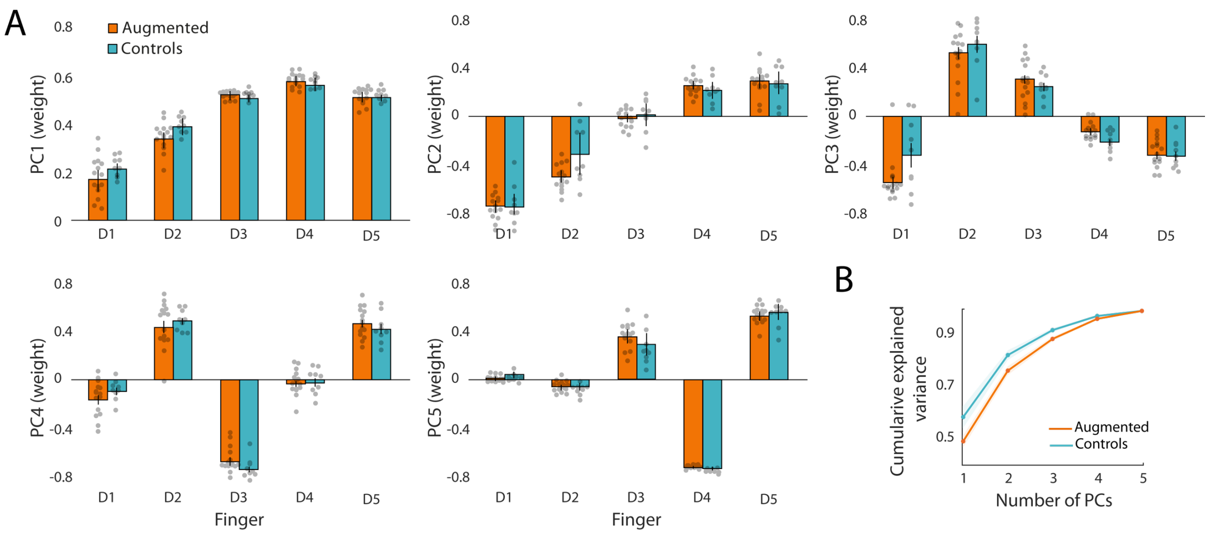


**Supplementary Figure S3**. (A) Average kinematic synergies captured during Thumb use. The first principal component (PC) reflects coordinated flexion/extension of all digits, while remaining PCs correspond to more individualised patterns of finger cooperation. (B) In the augmentation group, more principal components are needed to explain the same amount of variance as in the control group.
